# Single-cell transcriptomic analyses define distinct peripheral B cell subsets and discrete development pathways

**DOI:** 10.1101/2020.09.03.281527

**Authors:** Alexander Stewart, Joseph Ng, Gillian Wallis, Vasiliki Tsioligka, Franca Fraternali, Deborah Dunn-Walters

## Abstract

Separation of B cells into different subsets has been useful to understand their different functions in various immune scenarios. In some instances, the subsets defined by phenotypic FACS separation are relatively homogeneous and so establishing the functions associated with them is straightforward. Other subsets, such as the “Double negative” (DN, CD19+CD27-IgD-) population, are more complex with reports of differing functionality which could indicate a heterogeneous population. Recent advances in single-cell techniques enable an alternative route to characterise cells based on their transcriptome. To maximise immunological insight, we need to match prior data from phenotype-based studies with the finer granularity of the single-cell transcriptomic signatures. We also need to be able to define meaningful B cell subsets from single cell analyses performed on PBMCs, where the relative paucity of a B cell signature means that defining B cell subsets within the whole is challenging. Here we provide a reference single-cell dataset based on phenotypically sorted B cells and an unbiased procedure to better classify functional B cell subsets in the peripheral blood, particularly useful in establishing a baseline cellular landscape and in extracting significant changes with respect to this baseline from single-cell datasets. We find 10 different clusters of B cells and applied a novel, geometry-inspired, method to RNA velocity estimates in order to evaluate the dynamical transitions between B cell clusters. This indicated the presence of two main developmental branches of memory B cells. One involves IgM memory cells and two DN subpopulations, culminating in a population thought to be associated with Age related B cells and the extrafollicular response. The other branch involves a third DN cluster which appears to be a precursor of classical memory cells. In addition, we identify a novel DN4 population, which is IgE rich and on its own developmental branch but with links to the classical memory branch.

## Introduction

B cells can differentiate into plasma cells and secrete large amounts of antibody. There are, however, many more B cell functions which contribute to an effective immune response. B cells are highly effective antigen presenting cells, capable of presenting both protein and lipid antigens to T and NKT cells. B cell – T cell interactions trigger a variety of activation signals in both directions resulting in enduring affinity matured B cell memory, that may also be class switched, and activated T helper cells. B cells can also be activated via TLRs, producing pro-inflammatory cytokines such as IL6, TNFα and IFNγ in the process and resulting in differentiation into short-lived plasmablasts. The former, T-dependent, response will involve formation of germinal centres over time and, since it is dependent on T cells for maturation which have also been through tolerance checkpoints, it would normally have low risk of producing autoantibodies. The latter, extrafollicular, B cell response has the advantage of being more rapid, but also runs some risk of producing lower specificity antibodies. B cells can also be regulatory, producing IL10 and ensuring that autoreactive responses are not perpetuated.

In studying different functions of human B cells in health and disease most studies rely upon phenotypic differentiation in FACS analyses using IgD and CD27, or CD24 and CD38, in conjunction with the pan B cell marker CD19. For example, the CD19+CD27+IgD+ IgM memory population (1,2) is reduced in the elderly as a percentage of total B cells (3,4) This has important consequences for older people, since the IgM memory population is thought to provide protection against the bacterial polysaccharide T-independent antigens. Higher dimensional phenotyping shows that the IgM memory population in the blood is heterogeneous and further age-related differences are also seen (5), although the likely functional significance of this age related heterogeneity has yet to be determined.

The CD19+IgD−CD27− ‘double negative’ (DN) cells are of particular interest. Many different roles have been ascribed to this population; ‘memory precursors’, “exhausted memory cells”, ‘tissue based memory’, ‘extrafollicular ASC precursors’ and ‘atypical memory’ (6–15) or the most recent nomenclature DN1 (memory precursor) and DN2 (extrafollicular ASC precursor B cells) (16). DN cells are increased in older people, and in chronic infections such as HIV (6,7,10,11). DN cells are also expanded in autoimmune disease such as Systemic Lupus Erythematosus (SLE) (14,17) where they are responsive to IFNγ and thought to be precursors for pathogenic antibody secreting cells (8,9,18). Repertoire studies to try and clarify the relationship of DN cells to classical CD19+CD27+IgD− memory cells have been carried out and find both evidence for a close relationship with classical cells, with clones shared in both populations, as well as a difference in overall average repertoire character, with less hypermutation and larger complementarity determining regions (19). It is therefore likely that the DN compartment is functionally heterogeneous and only with high resolution techniques, such as single-cell transcriptomics, will it be possible to tease apart the sub-populations.

Single-cell transcriptomics is rapidly becoming a key methodology in biology thanks to its high resolution in terms of individual cells and high dimensional data. It offers the ability to discriminate between subsets of heterogeneous populations to understand individual contributions which may have previously been confounded by the Simpson’s paradox of studying averaged data. Unsupervised clustering algorithms offer us the chance to define subsets transcriptionally and interrogate the results to find tractable markers for use in phenotypical distinction of the same. Information about the possible functions of cell clusters can be inferred from the transcriptome relative to other clusters. scRNAseq is particularly useful in B cell immunology, where it has made pairing of heavy and light chain sequences possible (20).

Curated reference databases, such as the Human Cell Atlas (21), are important for comparison to experimental datasets. A frequently occurring problem in single-cell datasets is the relative lack of B cells, given that most such experiments are run on PBMCs; where B cells only make up between 5-10% of the total. Hence B cells are often under-described in single-cell experiments and often only divided into naive, memory and plasma cells which may miss important distinctions in disease. This is a general problem but has been particularly evident in recent Covid19 papers (16,22–28). We have therefore produced a dataset looking exclusively at five commonly defined B cell populations by phenotype to improve the resolution and better understand the heterogeneity present in B cells. By producing this dataset, we reconcile known phenotypic populations with those found in the transcriptome and can identify previously unknown populations in high dimensional space, thus providing a basic map of peripheral blood B cells as a reference dataset for the B cell community. We describe 10 major populations of B cells, including a novel population of IgE-expressing cells in the double negative compartment.

## Materials and Methods

### Samples, library preparation and sequencing

Peripheral blood mononuclear cells were isolated from a male healthy volunteer aged 25 using BD-bioscience lithium heparin vacutainers™ and Ficoll-Paque Plus (GE-healthcare). Cells were stained in BD Horizon Brilliant stain buffer with (Biolegend: CD19 HIB19 BV421, CD27 M-T271 FITC, IgD AI6-2 BV785, CD10 HI10a BV605, Live/Dead Zombie NIR), doublet and dead cells removed and lymphocytes gated on FSC/SSC, and sorted into Transitional (CD19+IgD+CD27−CD10+), Naïve (CD19+IgD+CD27−CD10−), IgM Memory (CD19+IgD+CD27+), Classical Memory (CD19+IgD−CD27+) and Double Negative (CD19+IgD−CD27−) populations on a FACSAria (BD Biosciences) at 4°C (Supplementary Figure S1). Cells were washed twice (5 min at 400 g, supernatant removed and replaced) in PBS supplemented with 0.04% non-acetylated BSA with a final spin through a 40 uM cell strainer. Populations were counted and run on a Chromium 10X controller using 5’ chemistry (10X Genomics) in individual lanes with an expected recovery rate of 4,000 cells per lane, according to the manufacturer’s instructions. Libraries were generated according to 10X genomics instructions and run on a High Output HiSeq2500 at 1 library per lane in 30-10-100 format.

### Data preprocessing, clustering and differential expression

Data was processed through CellRanger (10X Genomics, v3.1.0) and aligned to the GRCh38 genome. The raw transcript count matrix was loaded into R (v3.6.3) using the Seurat (v3.0) package (29,30). Cells were selected for further analyses according to the following criteria: (i) express zero *CD3E*, *GNLY*, *CD14*, *FCER1A*, *GCGR3A*, *LYZ*, *PPBP* and *CD8A* transcripts, to exclude any non-B cells and; (ii) express at least 200 distinct genes. Additionally, cells with total transcript count in the top 1% percentile were removed, as these cells were manually inspected to express transcripts of multiple V gene families per cell, indicating possible cell clumps tagged with the same barcode. In total, we have considered 8,771 cells and 16,093 genes. Data was log-normalised and the top 2,000 variably expressed genes were extracted using a variance-stabilising transformation (vst) as implemented in the FindVariableFeatures function in Seurat. The following genes were removed from the list of variably expressed genes in order to prevent downstream dimensionality reduction and clustering to reflect individual/clonotype specific gene usage: all Ig V, D, J genes (extracted using the regular expression [regex] “*IG[HKL][VDJ]*”), Ig constant genes (*IGHM, IGHD, IGHE, IGHA[1-2], IGHG[1-4], IGKC, IGLC[1-7]*, and *AC233755.1* [which encodes IGHV4-38-2]), *IGLL* genes, T-cell receptor genes (regex “*TR[ABGD][CV]”*). This pruned the variably expressed gene list to *n* = 1,840 genes. Principal Component Analysis (PCA) was then performed on this pruned gene list. Surveying the first 50 principal components, the proportion of variance explained plateaued at ~ 1.5% from the 15^th^ PC onwards. Uniform Manifold Approximation and Projection (UMAP) was performed based on the first 14 PCs, using the implementation in the python umap-learn package with correlation as the distance measure in the PC space. UMAP projections were produced on both two-dimensional and three-dimensional spaces. To define cell clusters, a shared nearest neighbour (SNN) graph was constructed using Seurat::FindNeighbors based on the first 14 PCs, and cell clusters were defined on the SNN graph with Seurat::FindClusters (resolution parameter = 1). Clusters were named according to manual inspection for their composition in terms of the original FACS-defined populations. Differential expression was examined using the Wilcoxon rank-sum test provided in Seurat::FindMarkers.

### Trajectory and RNA velocity analysis

To study the trajectory across the Seurat-defined cell subsets, a spanning tree across the data points was inferred using the monocle3 package (v0.2.0), based on the 3D UMAP embedding produced as detailed above. To estimate RNA velocity, spliced and unspliced transcripts were enumerated using the velocyto package (v0.17) (31). RNA velocity stream was mapped onto the UMAP space according to the pipeline provided in the SeuratWrappers package (v0.1.0) which uses functionalities provided in the velocyto.R package (v0.6) (31).

Based on the inferred velocity stream we also scored the transition between cell clusters using a geometrical approach. We evaluated, for each arrow depicting the velocity stream, its alignment with the projection towards cluster centroids, using the following procedure (illustrated in Supplementary Figure S2): first, for each arrow *A*_*i*_ with starting point *X*_*i*_ and ending point *Y*_*i*_, we determined the ‘starting cluster’ (i.e. the original cell identity from which transition is considered) by a majority vote of the identities of cells within the grid around *X*_*i*_ as given when the velocity stream was overlaid. We next considered, for every ‘destination’ cell cluster, the distance between *X*_*i*_ and its cluster centroid *Z*_*j*_. This permitted calculation of the angle q between and *X*_*i*_*Z*_*j*_, using the cosine formula:

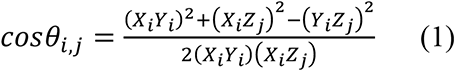

for each arrow *I* and each ‘destination’ cluster *j*. q should approach zero (and hence cos*θ* = 1) if the arrow is perfectly aligned to the direction pointing from the starting cluster to the destination. This forms the basis to define a ‘transition score’, taking into effect of the strength of the velocity (represented by the size of the arrow projected – i.e. *X*_*i*_*Y*_*i*_; it should positively contribute to the score) and the distance *X*_*i*_*Z*_*j*_ (which should negatively impact the score). This score, here denoted S, is given by:

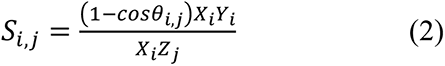

For every *I* and *j* the (1−cos*θ*) functional form ensures *S*_*i,j*_ increases as the arrow becomes more geometrically align to point to cluster *j*. This was performed for every arrow *I* and every cluster *j*. To summarise scores from individual *i*’s into a composite transition score (CTS) between every starting cluster *j*_1_ and destination cluster *j*_2_, all *i*’s are classified by their starting cluster and *S*_*i,j*_ are summed together for each *j*. Finally the CTS, i.e.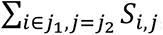, are adjusted by a weighting factor based on the distance between cluster centroids *d*_1_ (for cluster *j*_1_) and *d*_2_ (for *j*_2_). We first computed the pairwise distance between *d*_1_ and *d*_2_. Then, for each *d*_2_ (which corresponds to the destination cluster), we scaled all distances into a weighting factor *w* of range [0,1]:

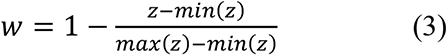

for all distances *z* pointing towards *d*_2_. Again, this functional form implies heavier weight is given for clusters which are close to one another in the dimensionality-reduced space. The final CTS was scaled by this factor:

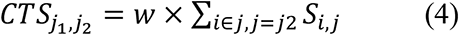

For our datasets RNA velocity streams were considered on two perspectives to view the 3D UMAP space: (1) UMAP1 vs UMAP3 (2) UMAP2 vs UMAP3, as visually these two perspectives separate the clusters the best. All arrows visualised in these two perspectives contributed towards the final CTS. These were plotted in Figure 2f in log_2_ scale to dampen extreme signals.

### Protein-protein interaction network inference

The R package ACTIONet (v1.0) (32) was used to infer protein-protein interaction networks (PPINs) for (i) each FACS-defined population; (ii) each cell cluster defined by Seurat, and; (iii) cells positive for T-bet (*TBX21*) transcripts. ACTIONet takes as input a reference protein interactome and uses the transcriptomic data to filter and score protein interactions relevant to a given cell subset. Here, the Unified PPIN (UniPPIN), which we previously compiled and recently updated (April 2020; Marzuoli, Ng and Fraternali, manuscript in preparation), was used as the reference interactome. UniPPIN contains in total of 17,997 proteins and 647,508 interactions, after mapping UniProt identifiers to HGNC gene symbols using the biomaRt (v2.42.1) R package. To reduce dimensionality, genes expressed in less than 10 cells were omitted, leaving a total of *n* = 12,867 genes for PPIN inference. Raw gene counts were used to calculate a ‘gene specificity’ score using the compute.annotations.feature.specificity function in ACTIONet, for each cell subset as defined above. This step enhances signals from genes exhibiting expression patterns specific to distinct cell subpopulations and penalises commonly-expressed genes. These gene specificities were subsequently used to infer a PPIN for each cell subset using ACTIONet::run.SCINET.annotations. Each node (i.e. protein) in the network is scored by a ‘specificity’ metric, which quantifies the centrality of the protein in the network scaled by its gene expression specificity. Since ribosomal and mitochondrial proteins frequently formed cliques (highly interconnected components) in biological networks, in order to mediate the impact of the inclusion of these proteins in analysing the topological and functional properties of the inferred PPINs, all ribosomal and mitochondrial proteins were removed from all the inferred networks prior to any statistical analyses and functional annotations.

### Functional annotation

Gene sets from Gene Ontology (GO) biological processes (BP) were downloaded from MSigDB (v7.1). Cluster-specific markers were annotated for GO BPs, by testing for gene set overrepresentation using Fisher exact tests. Gene Set Enrichment Analysis (GSEA) was also carried out (using the fgsea package [v1.4.1]) on the ACTIONet-inferred PPINs, with the proteins ranked using the ACTIONet per-node specificity metric.

### Data visualisation

Reads coverage was visualised using the Integrated Genomics Viewer (IGV, v2.8.2). All statistics and data visualisation were performed under the R statistical computing environment (v3.6.3). Heatmaps of gene expression were produced using pheatmap (v1.0.12). Visualisations of PPINs were produced using visNetwork (v2.0.9). Volcano plots were produced using EnhancedVolcano (v1.4.0). Visualisation of three-dimensional data embeddings were performed using the rgl package (v0.100.54). All other data visualisations were produced using functionalities provided in Seurat and the ggplot2 package (v3.3.0).

## Results

### B cell subsets

B cell sorted single-cell transcriptome libraries from five FACS sorted populations based on IgD/CD27/CD10 (Transitional ‘Trans’, Naive, IgM Memory ‘M-mem’, Classical Memory ‘C-mem’ and Double Negative ‘DN’) clustered into 10 unique populations using UMAP (Figure 1a). Of the 10 clusters formed in the UMAP analysis in all but 1 case (DN3) over 75% of the cells matched with their original FACS sorted population (Figure 1b). Differential gene expression analysis highlights these discrete populations (Figure 1c). Even though these cells were CD19 sorted only 34.4% percent of cells were CD19+ in the transcriptome, CD20 proving to be a much more reliable B cell marker in the transcriptome (Figure 1d)(29). Similarly, CD27 mRNA was not as abundant as CD27 surface protein, particularly in the C-mem cells. Importantly, we have explicitly filtered any immunoglobulin-related genes prior to clustering to avoid biasing the clustering algorithms by light chain or class of antibody. Even so, we find that *IGHE* and *IGHA2* cells can be differentiated by their transcriptome: DN3 being enriched for *IGHA2* and DN4 for *IGHE*. After finding the unusual DN4 IgE population, further investigation leads us to postulate that it is due to a cat hair allergy, exposure 3 days prior to sampling. We do not see this high level of *IGHE* in other data (unpublished) from peripheral blood. M-mem1 cells were distinguished from M-mem2 by higher IgD expression (Figure 1d). Transitional and Naive cells are best distinguished from other B cells by expression of *TCL1A*, higher in transitional, present in Naive and absent in all other cells.

**Figure 1.**
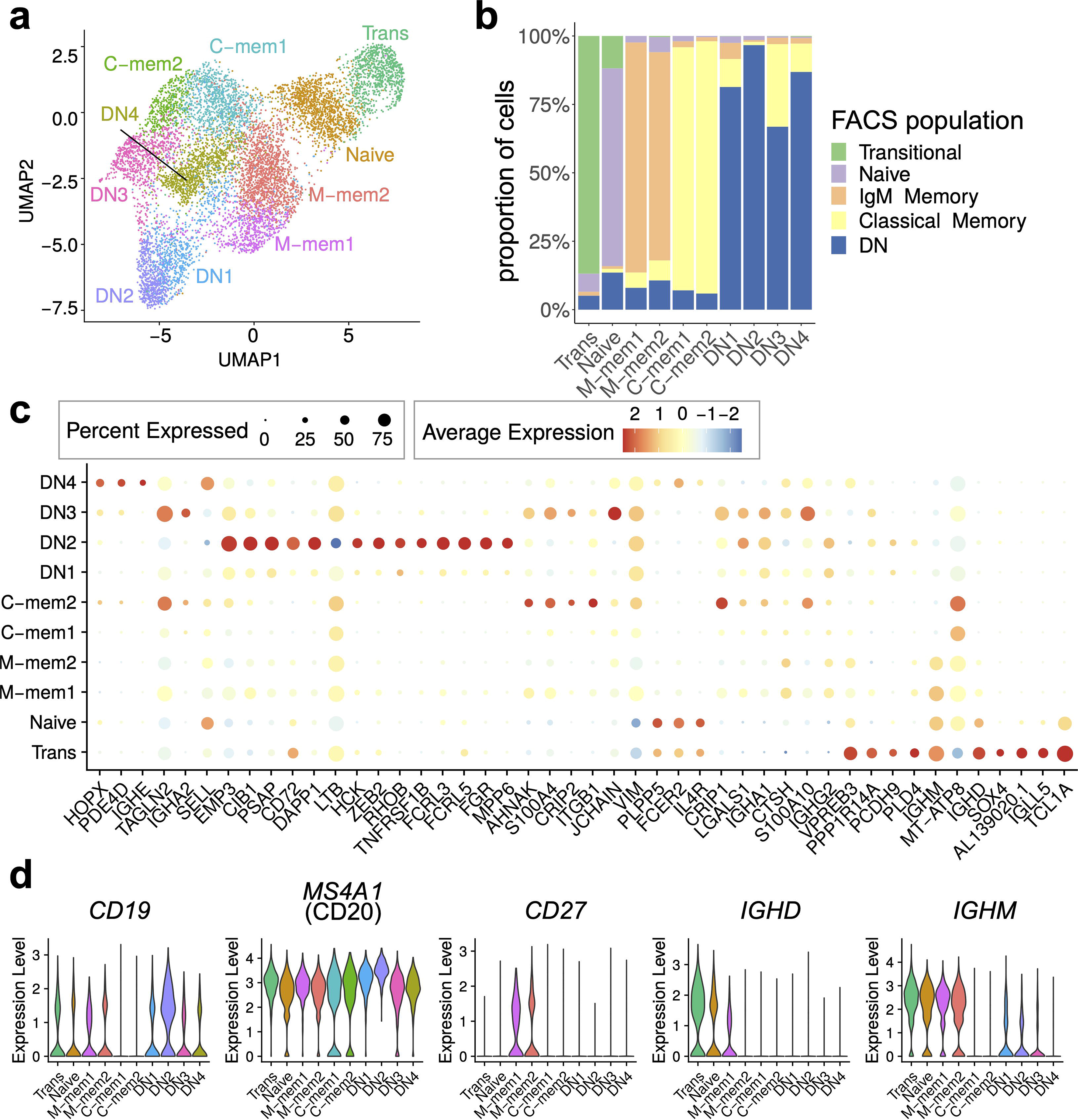
Single-cell atlas of peripheral B cell subsets for one individual. (a) Two dimensional UMAP projection of scRNA-seq data of peripheral B cell subsets. (b) Breakdown of each cell cluster defined using scRNA-seq data, in terms of the FACS-defined B cell identities. (c) Expression of top markers for each cell cluster. Here only markers with average log fold difference > 0.75 for at least one cluster are included. (d) Expression of *CD19*, *MS4A1* (CD20), *CD27*, *IGHD* and *IGHM* across cell clusters.

### Relationships between B cell subsets

We performed dimensionality reduction and pseudotime analyses to decipher the relationships between clusters and RNA velocity measurements were added to superimpose directionality information onto the trajectory (Figure 2). Four distinct branches of B cell clusters were seen in three dimensions (Figure 2a); 1) a transitional and naïve branch, 2) a classical memory and DN3 branch, 3) an IgM and DN1/2 branch and 4) a separate branch for the IgE-high DN4s (Figure 2a,b). In branch 1 the clear direction of development was transitional to naïve to memory and markers such as *CD38* and *TCL1A* are lost in the progression (Figure 2c). The relationships between the different memory populations are more complex (Figure 2a/d/e). We therefore developed a new method, by performing geometry-based calculations on the mapped velocity landscape (Supplementary Figure S2; also see Materials and Methods), to summarise the directionality, strength and position of individual RNA velocity streams between individual cell clusters. This produced a composite transition score (see Materials and Methods) that quantifies the developmental flow of cell clusters (Figure 2f) which can be mapped conceptually onto the inferred trajectory (Figure 2g). From this the M-mem2 population almost appears as a separate originating singularity with flow out of the IgM memory cluster, to DN1/2 and C-mem1, but with relatively little flow from Naïve into the M-mem clusters themselves (Figure 2 f/g). The unidirectional flow from DN1 to termination at DN2 is strong. We also see a very strong flow from DN3/4 towards C-mem1/2 terminating in C-mem2.

**Figure 2.**
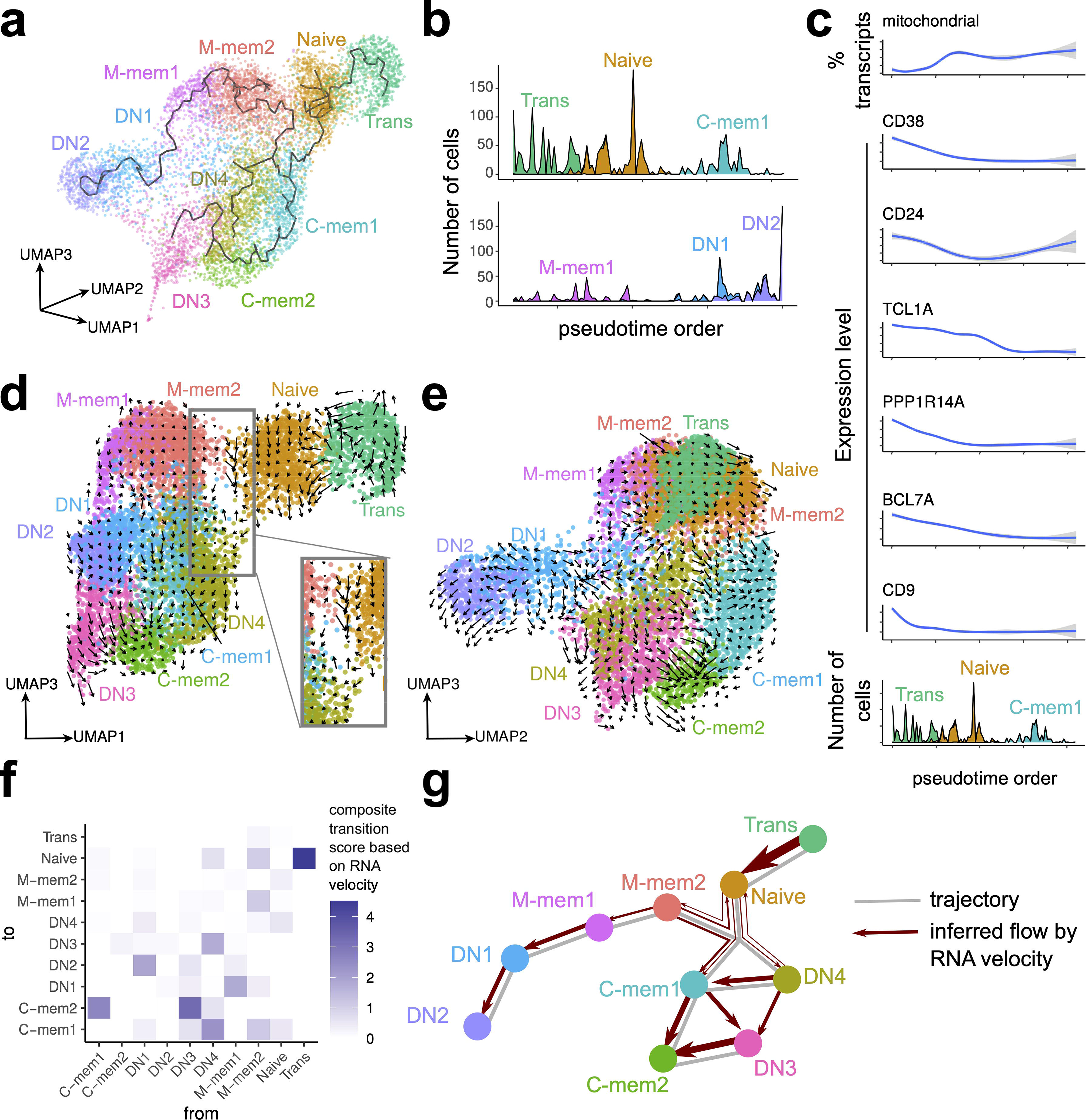
Trajectory and RNA velocity analyses of B cell clusters. (a) Trajectory inferred using monocle3, overlaid onto cell clusters in a three-dimensional UMAP space. (b) Pseudotime order of cells inferred using monocle3, in (top) Trans, Naïve and C-mem1, and (bottom) M-mem1, DN1, DN2 clusters. (c) Change in expression level of selected genes across the pseudotime axis for Trans, Naïve and C-mem1. (d-e) RNA velocity stream overlaid on three-dimensional UMAP space. Notice the different UMAP dimensions depicted in each panel. In (d) the inset represents a close-up view into the space between Naive, M-mem2 and the class-switched memory clusters. (f) Composite transition score between cell clusters, calculated based on examining geometrically the alignment of velocity streams to cluster positions. (g) Summary of trajectory and RNA velocity analyses.

A UMAP projection considering only the DN population highlights the distinct and robust partitioning of these cells into 4 subpopulations (Figure 3a). DN1 and DN2 display similar immunoglobulin class distribution, although DN1 has more IgM and DN2 slightly higher IgG2/3 expression. DN3 is IgA2-rich, while DN4 is IgE-rich (Figure 3b).

**Figure 3.**
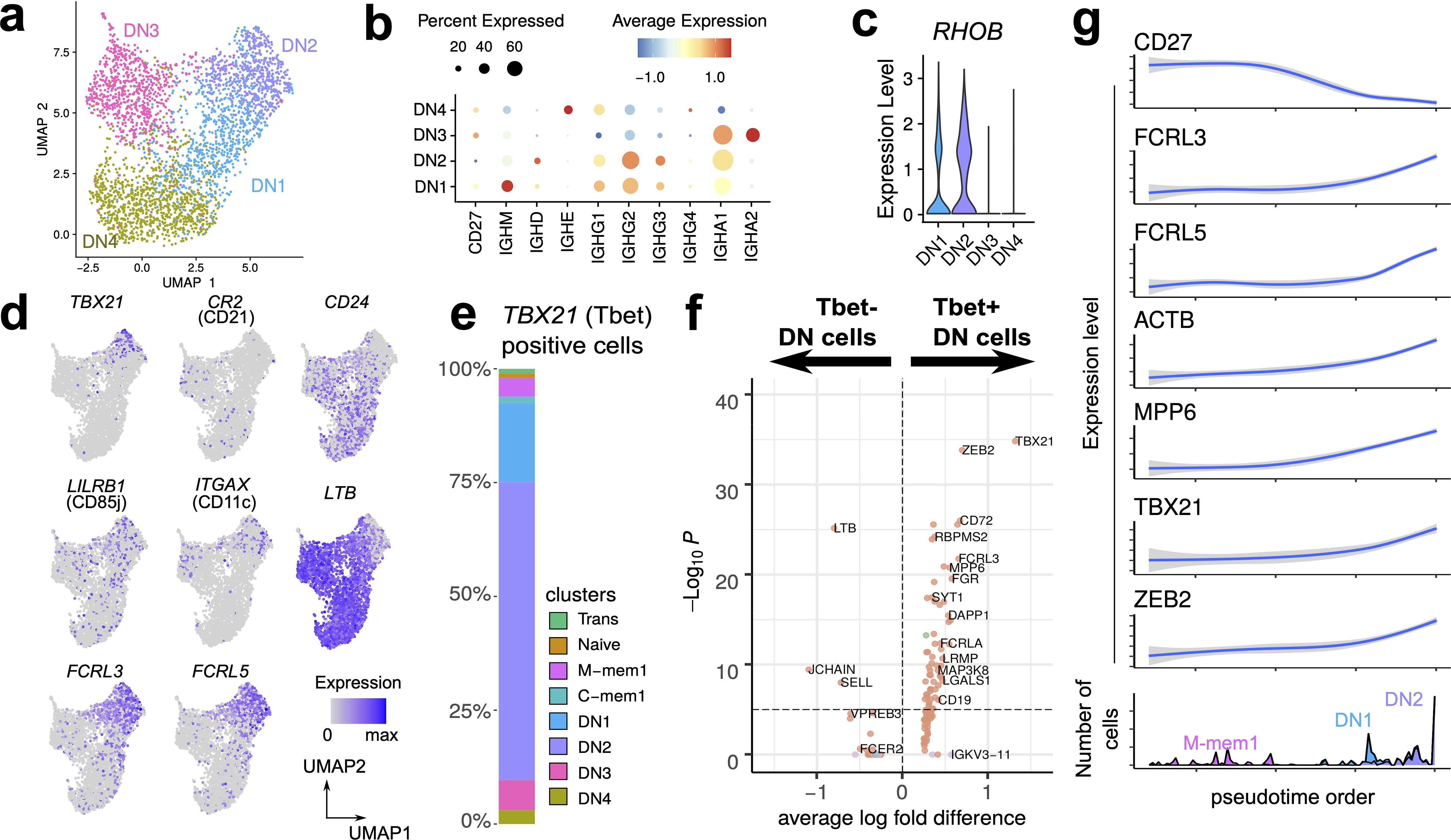
Heterogeneous single-cell expression landscape of double-negative memory B cells. (a) Dimensionality projection of DN1-4 clusters on a two-dimensional UMAP space. (b) Expression of *CD27* and *IGH* constant region genes across the four DN clusters. (c) Expression level of *RHOB* across the four DN clusters. (d) Expression of positive and negative markers of DN1 and DN2 clusters. (e) Breakdown of cells positive for *TBX21* transcripts (which encodes Tbet) by the cell clusters defined in this dataset. (f) Differential expression analysis of Tbet+ and Tbet-DN cells. (g) Change in expression levels of selected genes through the pseudotime order of M-mem1, DN1 and DN2 cells.

As a unifying marker both DN1 and DN2 have higher expression of *RHOB* (Ras Homolog Family Member B, involved in intracellular protein trafficking) compared to DN3 and DN4 (Figure 3c). The DN2 population at the terminal point of the Mmem/DN development branch is of interest, expressing T-bet (*TBX21*), CD11c (*ITGAX*), and lacking *CD21* (Figure 3d, Supplementary Figure 3) concomitant with an identity of Age-related B cells (33). This population is also enriched in the inhibitory receptors *FCRL3*, *FCRL5* and lacks lymphotoxin-B (*LTB*), *CD24* and *CXCR5* (Figure 3d). These are features in common with a previously identified precursor population for extrafollicular antibody secreting cells (ASCs) (16). T-bet has been shown to mediate IFNγ-dependent (but not IL4 dependent) development of antibody secreting cells from B cells (34). With this in mind we looked at all cells that were T-bet positive and showed that the large majority were in the DN population in the order DN2>DN1>DN3>DN4 (Figure 3e). Other transcriptional markers for the T-bet+ cells include *ACTB*, *MPP6*, (Figure 3 f,g) and *ALOX5AP*, *GSTP1*,*LAPTM5* (Figure 4a)

**Figure 4.**
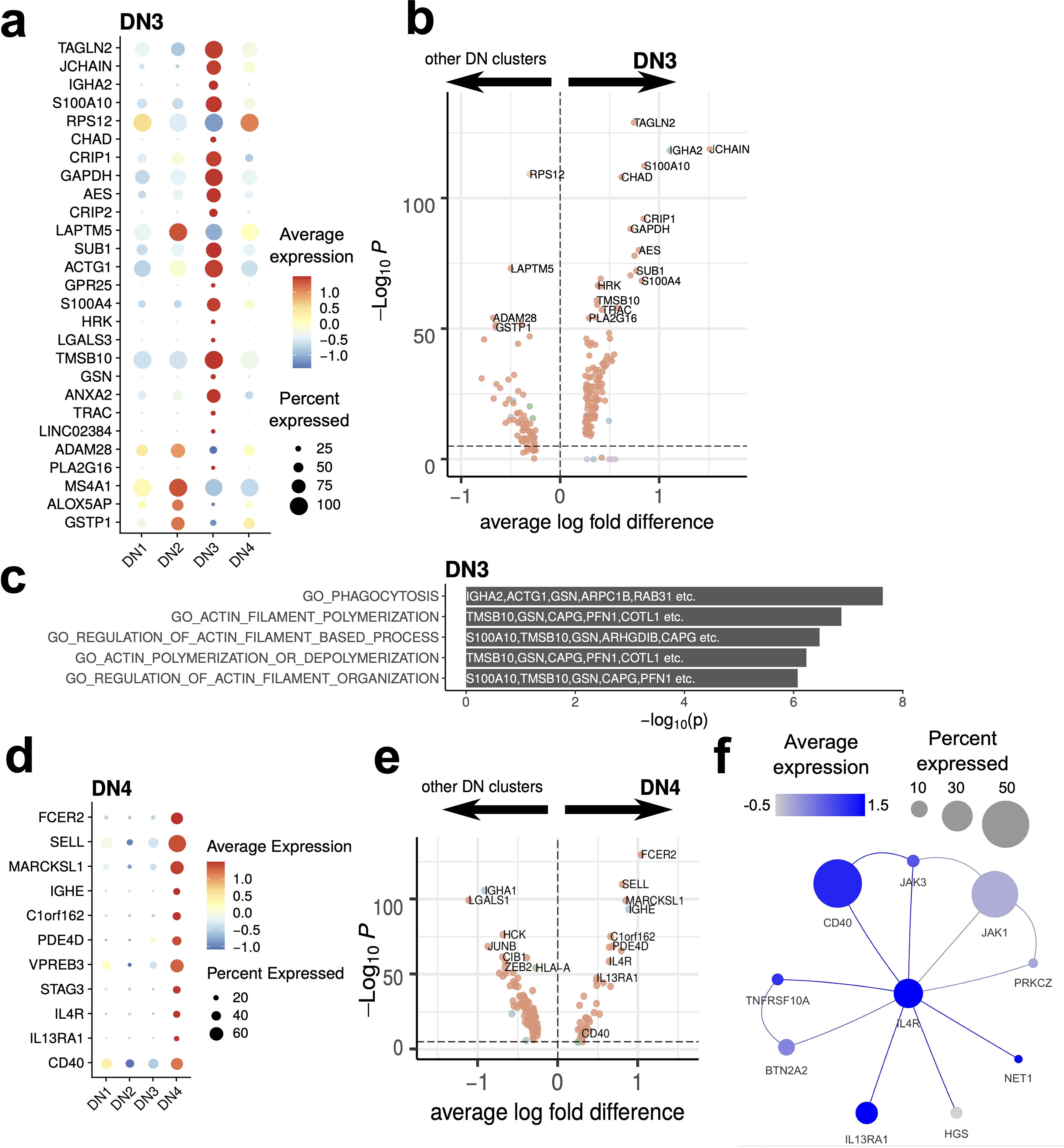
DN3 (IgA2-high) and DN4 (IgE-high) DN cells. (a) Expression level of DN3 markers illustrated with a dot plot. (b) Differential expression analysis of DN3 compared with other DN clusters. (c) Gene ontologies of DN3 markers. The top 5 pathways in this GO overrepresentation analysis were illustrated. Top markers overlapping these pathways are noted on the plot. (d) Expression levels of DN4 markers illustrated with a dot plot. (e) Differential expression analysis of DN4 compared with other DN clusters. (f) Inferred protein-protein interaction network of *IL4R* and its direct neighbours. The expression of these genes in the DN4 cluster were mapped onto the nodes of the network.

The DN3 population expresses high levels of *IGHA2* and *JCHAIN* (Figure 4). Upregulation of Transgelin-2 (*TAGLN2*) expression suggests an activated state (35). Inferred protein-protein interaction network (PPIN) analysis of the DN3 population shows genes enriched in proteins involved in actin filament formation and organisation (Figure 4c). This population falls on a separate differentiation pathway from DN1 and DN2 and more closely resembles classical Memory B cells in differential gene expression (Figures 1c, 2g).

The IgE-expressing DN4 population has very high levels of *FCER2* expression, the low affinity receptor for IgE, as well as *IL4R*, *IL13RA1*, and the co-stimulator *CD40* (Figures 4d/e); all forming part of the same activation network (Figure 4f). This IgE population is therefore in an active state and suggests the donor was undergoing an allergic response.

Traditionally human memory B cells were known as CD27+IgD− and we have designated these “classical memory”. The transcriptomic clustering reveals two different subpopulations in our data: C-mem1 and C-mem2. The main differences between the two are in expression of mitochondrial and ribosomal genes (Figure 5a). This might reflect a low overall gene expression by C-mem1, meaning ribosomal genes are over-represented (Figure 5a). Coupled with their relative positioning in the pseudotime/RNA velocity analysis (Figure 2g) this could indicate a more quiescent population awaiting activation for C-mem1, while C-mem2 are more active with higher active mitochondrial gene expression. Comparison of C-mem with DN indicates that DN cells have higher levels of CD20 (*MS4A1*) and HLA genes (Figure 5b). Analysis of individual genes and inferred PPINs (Figure 5c-d) shows genes predominantly involved in protein translation are expressed by the C-mem1, while actin networks involved in movement, division and immune synapse formation, form the top networks for C-mem2.

**Figure 5.**
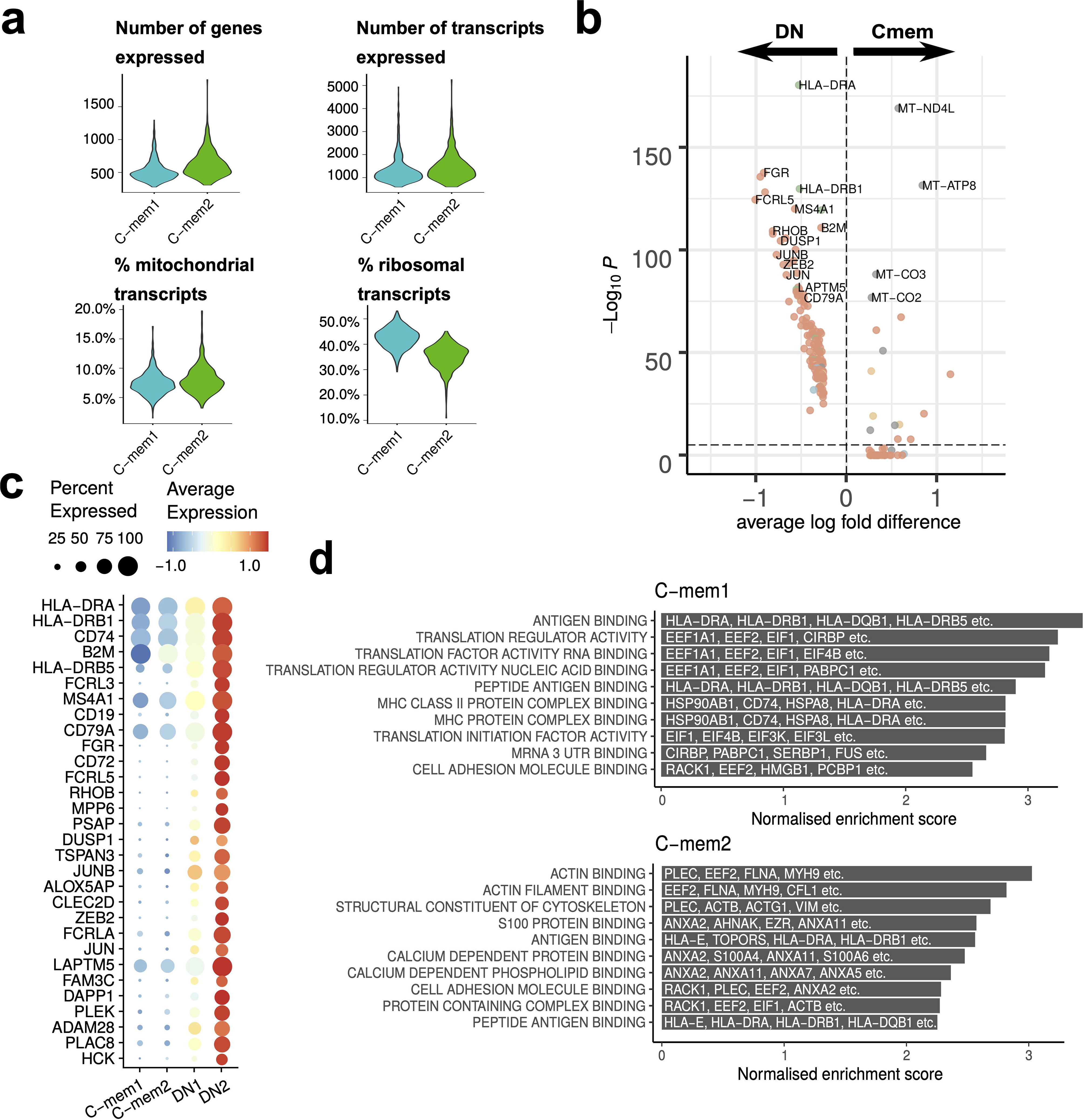
Comparison between classical memory (C-mem) and double-negative (DN) memory B cells. (a) Comparison of the number of genes and transcripts expressed, and the proportion of mitochondrial and ribosomal transcripts between the C-mem1 and C-mem2 clusters. (b) Differential expression analysis of C-mem and DN cells. (c) Expression of differentially expressed genes between C-mem1/2 and DN1/2 visualised in a dot plot. (d) Gene set enrichment analysis of the inferred PPINs for the C-mem1 and C-mem2 cells. The top 10 significant pathways are shown here.

IgM memory cells display a number of genes of interest (Figure 6a) including *CD44* involved in cell to cell interactions and activation (36,37) along with *MARCKS* involved in actin cross linking (38), *TCF4* (encoding E2-2) which recognised the E-box binding site originally identified in immunoglobulin enhancement (39) and *SMARCB1* which relieves repressive chromatin structure (40). Such a signature of expression would suggest an active group of cells involved in cell-to-cell interaction. We also highlight very strong expression of *CD1C*, a marginal zone B cell marker (41), in a small but distinct number of IgM memory cells.

**Figure 6.**
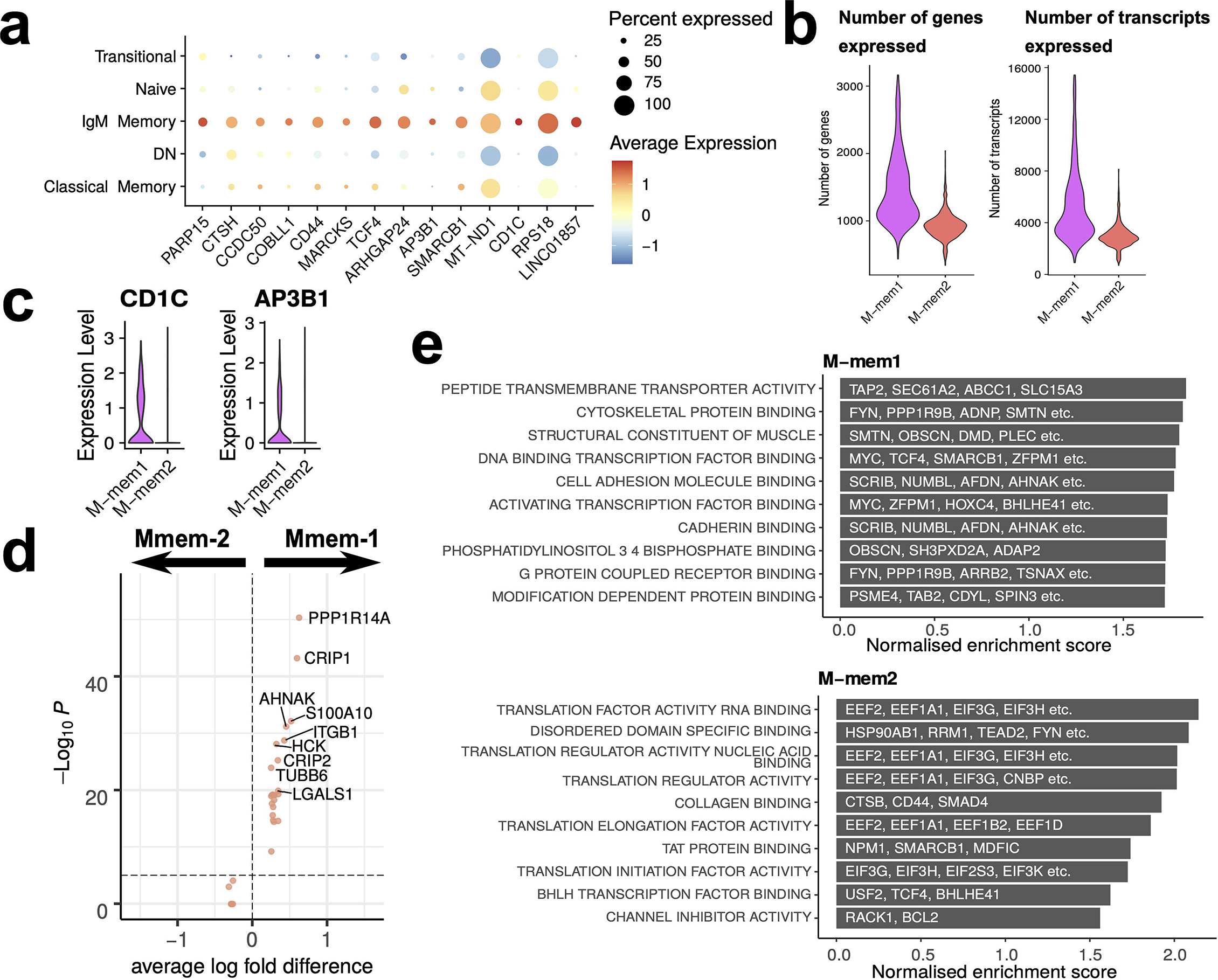
IgM memory cells. (a) Transcriptional markers for IgM Memory cells in comparison to other FACS-defined populations. (b) Numbers of genes and transcripts expressed per cell in M-mem1 and M-mem2 clusters. (c) Expression of *CD1C* and *AP3B1* in M-mem1 and M-mem2 clusters. (d) Expression of differentially expressed genes between M-mem1 and M-mem2 cells. (e) Gene set enrichment analysis of the inferred PPINs for the M-mem1 and M-mem2 cells. The top 10 significant pathways are shown here.

The two IgM memory subpopulations are predominately defined by the quantity of transcript rather than expression of any one set of genes, M-mem2 containing far fewer transcripts. Taken together with the position of these two subsets relative to one another in the pseudotime and velocity analysis, these suggest an activation pathway from a quiescent population M-mem2 to the active M-mem1 population. Most of the gene markers shown in Figure 6a are expressed in both the M-mem1 and M-mem2 clusters (Supplementary Figure S4) except *CD1C* and *AP3B1* which are uniquely expressed in M-mem1 cells (Figure 6c). Both genes are involved in presentation of lipid antigens (42). Very few other genes were differentially expressed between M-mem1 and M-mem2 (Figure 6d). The inferred protein interaction networks were more informative, showing translational housekeeping networks dominating the M-mem2 population while M-mem1 networks include both cell adhesion and structural networks suggestive of cell binding (Figure 6e).

## Discussion

The separation of B cells into functionally different populations is useful in trying to understand immune responses to challenge. Typically, this has been done using surface markers to separate cell types and then ascribing functions to the subsets. However, averaging effects over a heterogeneous population can quite often mask important information. The advent of single-cell technologies enables more precise differentiation between cell types and more understanding of their possible functions based on the genes transcribed. There is a wealth of immunological data based on phenotypic subset separation and maximisation of insight to be gained from all available information requires the unification of phenotype and transcriptome techniques. Here we used scRNAseq on the major peripheral B cell populations of an individual, FACS sorted by IgD, CD27 and CD10 (Transitional, Naive, IgM Memory, Classical Memory, Double Negative), to consolidate the phenotype with the transcriptome. Building upon conventional clustering, pseudotime and RNA velocity analyses, we devised a geometry-inspired method to summarise information from individual RNA velocity streams, and derived quantifications for the transitions between cell clusters (Figure 2f,g). RNA velocity analysis has been widely developed and applied since its conception (31,43), providing insights into the transcriptional dynamics of single cells. Individual velocity streams could be difficult to interpret, especially in systems where multi-way transitions amongst several cell clusters are possible. Here we devised a method to build upon these individual velocity streams and provide a summary score, which allowed us to consider the dynamical relationships between all the discovered cell clusters in an unbiased manner (Figure 2g).

We have found that the DN population (IgD−CD27−) clusters into four different sub-populations. The DN2 population differentially expressed more *TBX21* and *FCRL5* but no *CD24*, matching the descriptions of Age-related B cells and extrafollicular ASCs (8,14,16,33). Our DN1/3 populations both have increased levels of IgG/A genes and *CD24* with decreased *CD21* expression and they match previous descriptions of DN memory precursor populations (16,44). However, our data indicates that they are on different developmental trajectories. The RNA velocity data shows that DN3 cells flow predominately into the classical memory compartment, while DN1 cells flow predominantly into DN2 cells. DN3 may be related to a similar to a previously reported CD21−/CD11c− population (45,46). Together the transcriptome, UMAP analysis and RNA trajectories, all suggest that DN1 and DN3 are separate DN memory populations that contribute to two distinctly different branches of development with hardly any crossover between the two (Figure 2f,g). T-independent B cells responses are known to expand the IgM memory cell population and plasma cell lineage commitment (47–50) which would appear, on the evidence provided here, to be occurring through an IgM memory/DN1/DN2 pathway. We therefore suggest that these two pathways represent the T-independent (DN1) and T-dependent (DN3) B cell development pathways. The end points of both these pathways, DN2 and Cmem2, have RNA velocity arrows continuing to point outwards which may indicate further development, for example into ASCs.

IgM memory cells display several genes involved in actin regulation and cell to cell interactions that might suggest formation of B cell synapses. Expression of the gene *MARCKS* has recently been found to increase lateral motility of the BCR by modulating the cytoskeleton (38), CD44 is a known lymphocyte activator (36,37), *TCF4* (E2-2) a developmental transcription factor which regulates the IgG enhancer (23,39), and *SMARCB1* a chromatin re-modeller that relives repressive structures (51). The presence of a distinct number of IgM cells strongly expressing *CD1C*, marginal zone B cell markers (52) is supportive of the idea that these are circulating marginal zone cells following a T-independant pathway of differentiation (41,53,54). In contrast to other clusters, which have limitations as to their interactions with other groups, the RNA velocity data seems to suggest that cells from the Mmem2 cluster could give rise to cells in many different groups in all branches of the model (Figure 2f)

It is worth noting that the DN3 population is enriched in IgA, particularly genes encoding IgA2 and J chain. This, despite exclusion of immunoglobulin related genes from the clustering, indicates that IgA2-expressing cells are functionally different from the IgG-expressing cells (Figure 3b). In a similar fashion, we also found an IgE-expressing sub-population of DN cells, DN4. This population is over represented in the data due to our enrichment of the DN subset by FACS sorting and, in reality, represents approximately 0.5% of total B cells in this volunteer; DN making up 1.46% of this patients B cells (Supplementary Figure S1) and 34.21% of cells in the DN clusters are DN4. Even this level is unusually high and is assumed to be the result of exposure to cat hair 2 days prior to sampling although no symptoms were present at blood draw. CD27-IgE+ cells have been seen previously (55), alongside CD27+IgE+ cells, but in these data DN4 is the only cluster to show IgE expression (Figure 1c, 3b). The study of IgE-expressing cells is usually hampered by their scarcity, the poor polyadenylation of IgE transcripts and the difficulties in distinguishing IgE-expressing cells from those binding to IgE via Fc∊RII (56,57). Single cell transcriptomics can circumvent the antibody staining issues and shows here that this population seems to be activated, given the evident RNA velocity stream into and out of DN4 from other clusters (Figure 2g). The presence of *CD24* transcripts and RNA velocity flow into the classical memory population suggests these are memory precursors, although this is difficult to reconcile with the lack of IgE in the classical population. The tendency for some DN4 velocity arrows to point away from the UMAP plot might indicate that these cells are precursor to ASCs, as we have suggested for DN2 and Cmem2.

Both the C-mem and the M-mem sorted populations were split into two separate clusters. M-mem2 preceding M-mem1 and C-mem1 preceding C-mem2 in the RNA velocity plot. These separations appear to be largely due to transcriptional activity resulting in differential expression of ribosomal or mitochondrial genes. Such variability is possibly the result of a quiescent sub-population (C-mem1, M-mem2) rather than outright differences in function. In support of this, C-mem1 expresses networks dominantly related to antigen binding and housekeeping protein translation functions, but little else. In comparison, the three top networks of C-mem2 are actin and cytoskeleton related genes, suggestive of greater activity and regulation of the BCR leading cell to cell signalling (58).

We find that transitional and naive cell populations are transcriptionally homogenous. The T1, T2 and T3 transitional cells that have previously been distinguished by gradations of surface markers such as CD10, or CD24+CD38+, are not clustered separately in this data. RNA velocity highlights the highly directional development of transitional cells into naive cells, but only indicates small velocities out of the naive pool into memory. This may look different in an immune challenged volunteer where one would expect active differentiation between naïve and memory. However, the naïve to memory differentiation processes may occur in tissue and not be apparent in the blood.

The occasional disparities between mRNA and protein level gene expression means that markers in the transcriptome do not always translate into the proteome and *visa versa*. Our atlas of peripheral blood B cells can be used as a tool to identify the same cell subsets in other single-cell transcriptome datasets, using referenced-based bioinformatic approaches. It can also be used to identify new markers for tractable methods of B cell sorting (Supplementary Figure S5). It is worth noting that transcriptome data can suffer from lack of sequencing depth, so previously well-known phenotypic markers may not always be reliable at a single cell level. For example, CD27 suffers from incomplete transcript coverage. When clustering data from B cells we strongly suggest removal of variable genes and immunoglobulin constant regions as these tend to skew UMAP results. For example, the kappa lambda light chain distinction between cells is clear and the resulting skew can mask important transcriptomic differences between other, more functional, distinctions. To define naive and transitional cells we would suggest *TCL1A* to identify both populations and then *CD38*+*TCL1A*++ to identify transitional and *CD38*-to identify naive. IgM memory clusters can be identified as *TCL1A*-, *CD27*+, *IGHM*++, *IGHD*+. Markers for classical memory cells appear less reliable although can be used to broadly separate them. Although expressed in all cells, the classical memory cells also display higher mitochondrial gene expression than all other cell types. In cases where identifying memory cell populations is of importance we would advise using CD27 feature barcoding (also called CITEseq or TotalSeq) to overcome the low mRNA expression levels. Our sub-populations of quiescent Cmem and Mmem cells can only be distinguished by generally low gene expression and not by a defined set of markers. Identifying total DN cells is by *CD27*-, *IGHM*-, *TCL1A*-, *IGHD*- and the sub-populations separated as per Table 1. We also note that we only find *TBX21*+ cells in the DN population and not in the naive compartment as with SLE (8,9,59). Additionally, we did not find a population of *FCRL4*+ DN cells typical of HIV patients (60), highlighting the uniqueness of these populations to particular immune disorders.

**Table 1:**
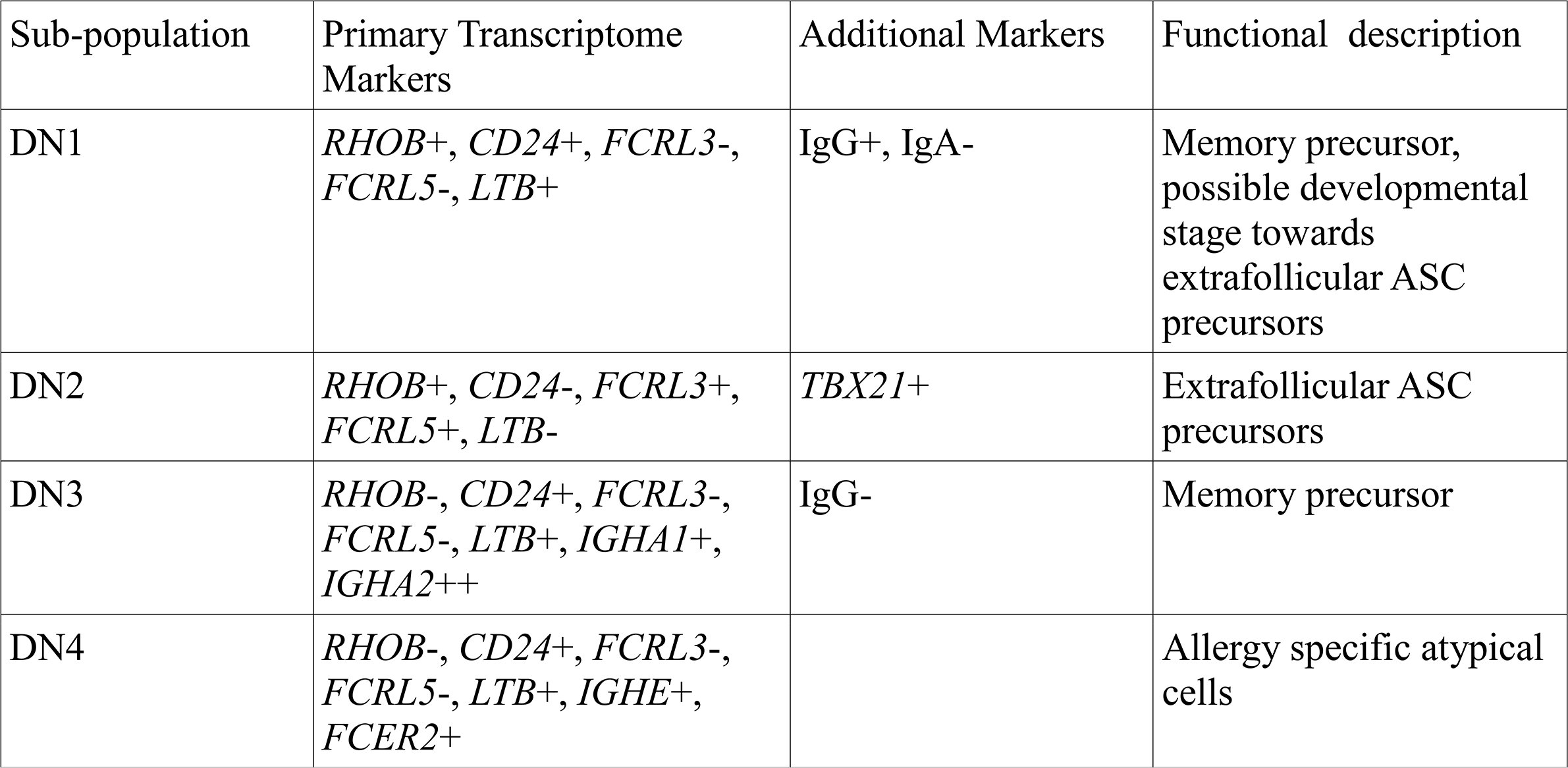
Classification of double negative sub-populations using transcriptome markers.

| Sub-population | Primary Transcriptome Markers | Additional Markers | Functional description |
| --- | --- | --- | --- |
| DN1 | *RHOB*<sup>+</sup>, *CD24*<sup>+</sup>, *FCRL3*<sup>-</sup>, *FCRL5*<sup>-</sup>, *LTB*<sup>+</sup> | IgG<sup>+</sup>, IgA<sup>-</sup> | Memory precursor, possible developmental stage towards extrafollicular ASC precursors |
| DN2 | *RHOB*<sup>+</sup>, *CD24*<sup>-</sup>, *FCRL3*<sup>+</sup>, *FCRL5*<sup>+</sup>, *LTB*<sup>-</sup> | *TBX21*<sup>+</sup> | Extrafollicular ASC precursors |
| DN3 | *RHOB*<sup>-</sup>, *CD24*<sup>+</sup>, *FCRL3*<sup>-</sup>, *FCRL5*<sup>-</sup>, *LTB*<sup>+</sup>, *IGHA1*<sup>+</sup>, *IGHA2*<sup>++</sup> | IgG<sup>-</sup> | Memory precursor |
| DN4 | *RHOB*<sup>-</sup>, *CD24*<sup>+</sup>, *FCRL3*<sup>-</sup>, *FCRL5*<sup>-</sup>, *LTB*<sup>+</sup>, *IGHE*<sup>+</sup>, *FCER2*<sup>+</sup> |  | Allergy specific atypical cells |

In summary, we show branching pathways of B cell development that appear to separate into T dependent and T-independent pathways. The subset of B cells previously known as “Double negative” contains a variety of different cells belonging to both pathways. DN1 and DN2 being in the T-independent pathway closely related to IgM Memory cells, and DN3 being more closely related to classical memory cells but enriched in genes encoding IgA2 and J chain. In addition, the serendipitous use of an allergic individual after allergen contact has helped to show an IgE-expressing B cell subset within the DN family, but clustering on its own separate branch of the RNA velocity pathway. That said, it seems to have strong links to the T-dependent development pathway containing classical memory cells and the DN3 cluster rather than the IgM memory/DN1/DN2 development pathway. The data we have collated can be used to map these B cell subsets in other single cell transcriptomic data.

## Data Availability

Raw sequencing reads, processed Seurat object, RNA velocity calculation and inferred interactomes were made available at ArrayExpress (accession E-MATB-9544). Code devised for calculating composite transition scores between cell clusters is available on github at https://github.com/Fraternalilab/scRNAVeloQuant.

## Acknowledgements

We thank the technical team at the University of Surrey for FACS support and the donor for their blood.

## Author Contributions

AS, FF, DDW devised and designed the experiment. AS carried out the experiment and data acquisition, with FACS sorting performed by GW and VT. JN and AS performed bioinformatics analysis. All authors contributed towards interpretation of data and writing of the manuscript.

