## Supplemental Figures for "Single-cell transcriptomic analyses define distinct peripheral B cell subsets and discrete development pathways"

**Supplementary Figures**

Supplementary Figure S1

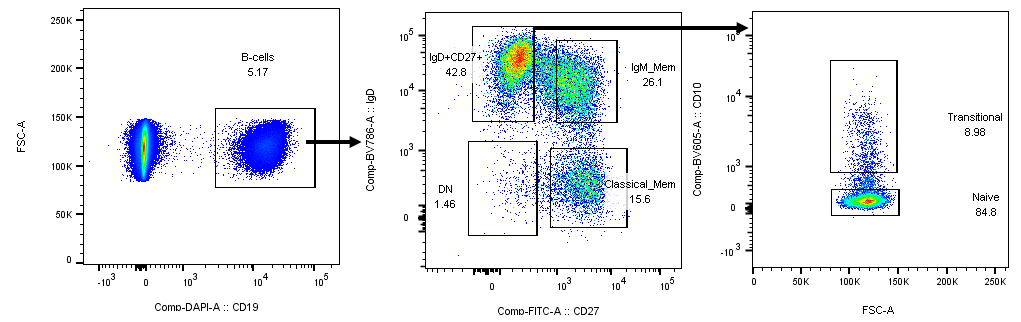

Gating strategies to sort B cell populations used in this work.

Supplementary Figure S2

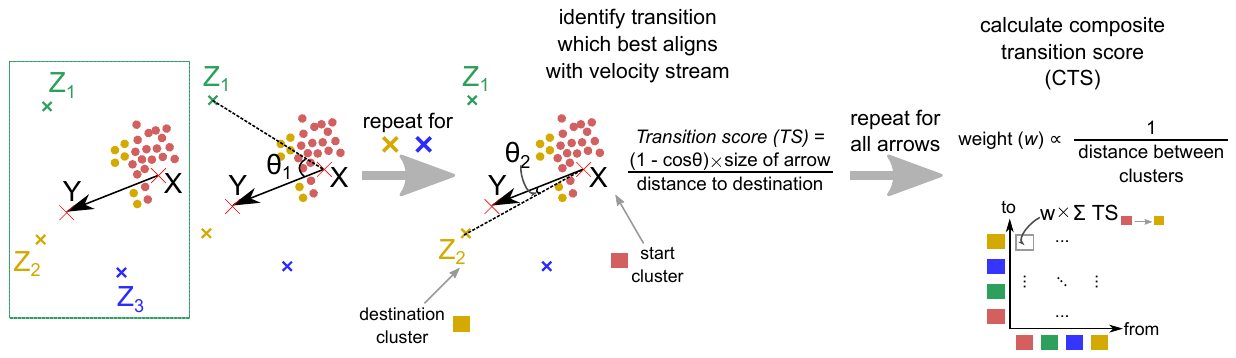

Calculating composite transition score between cell clusters using RNA velocity streams overlaid on the dimensionality-reduced data space. (*left box*) As an illustration a single velocity stream (indicated by the arrow XY) was considered in a space with four clusters, the starting cluster (here coloured red, represented by the red dots [i.e. cells] around point X) and three other clusters represented by their centroids Z_1_, Z_2_ and Z_3_. (*middle*) For each of Z_1_, Z_2_ and Z_3_, we compute the angle θ between XY and the projection XZ_i_. A transition score (TS) was calculated based on this geometric relation, and based on this the transition which best aligns with XY was identified (in this case to the yellow cluster with centroid Z_2_). (*right*) Considering all arrows overlaid on the data space, a composite transition score (CTS) is calculated to quantify the support in terms of RNA velocity for transition between any pair of clusters, weighted by their distances.

Supplementary Figure S3

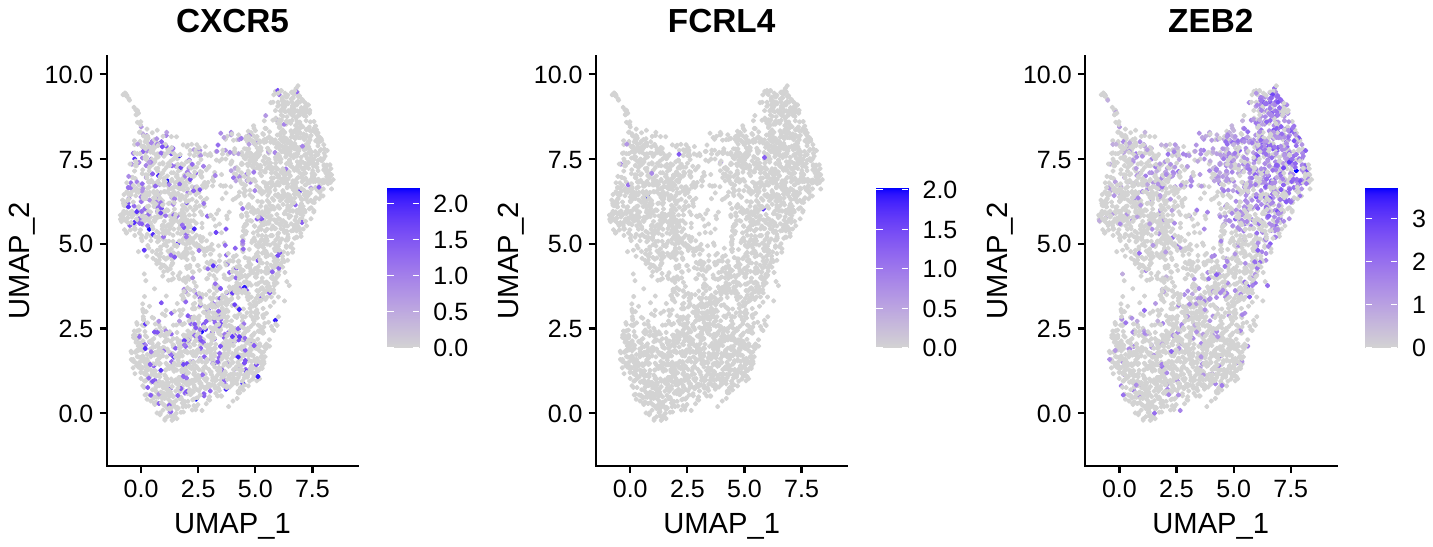

Expression of *CXCR5*, *FCRL4* and *ZEB2* across the four DN clusters.

Supplementary Figure S4

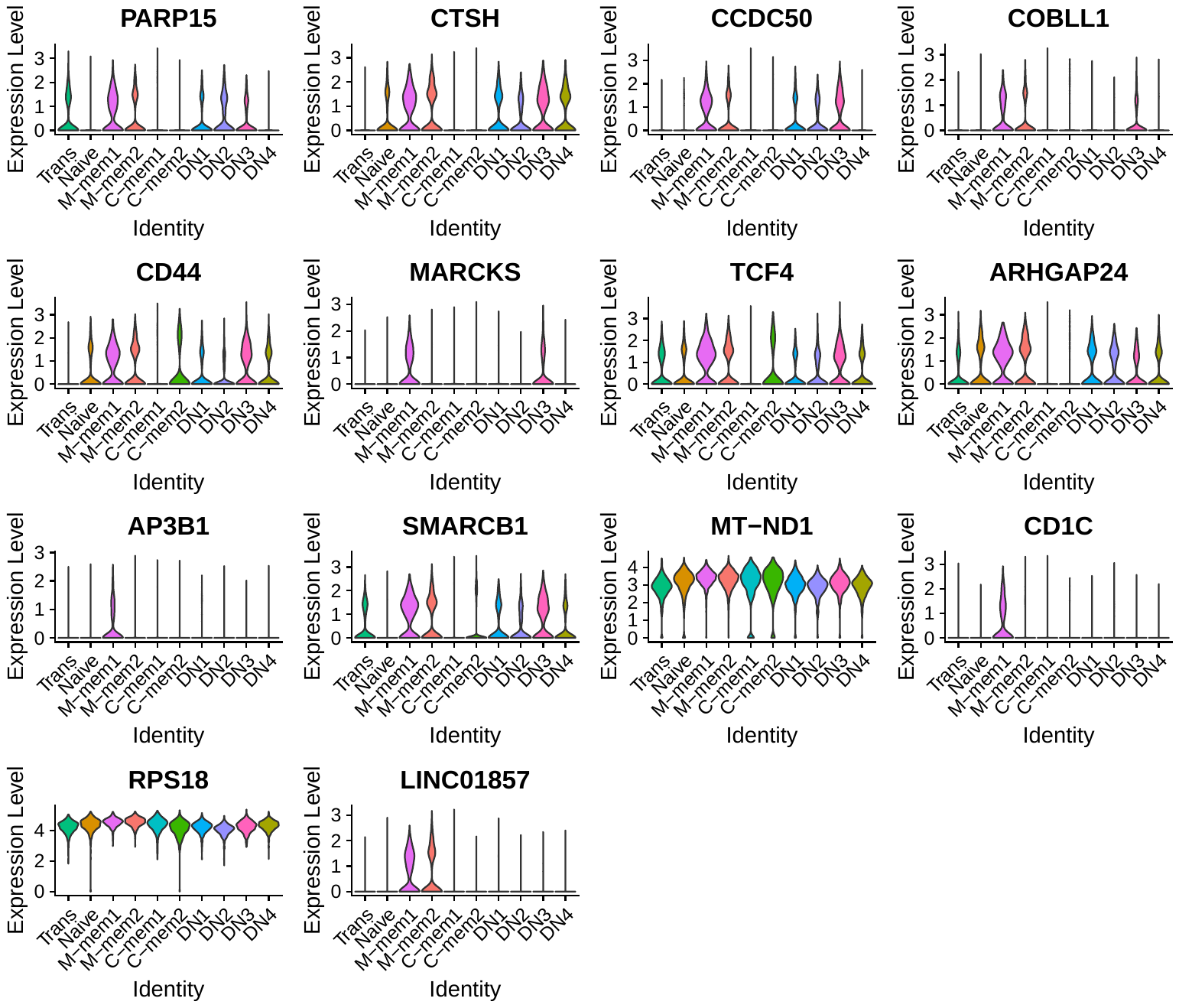

Expression of markers of IgM Memory cells shown in Figure 6A (main text), here depicted as violin plots across all cell clusters.

Supplementary Figure S5

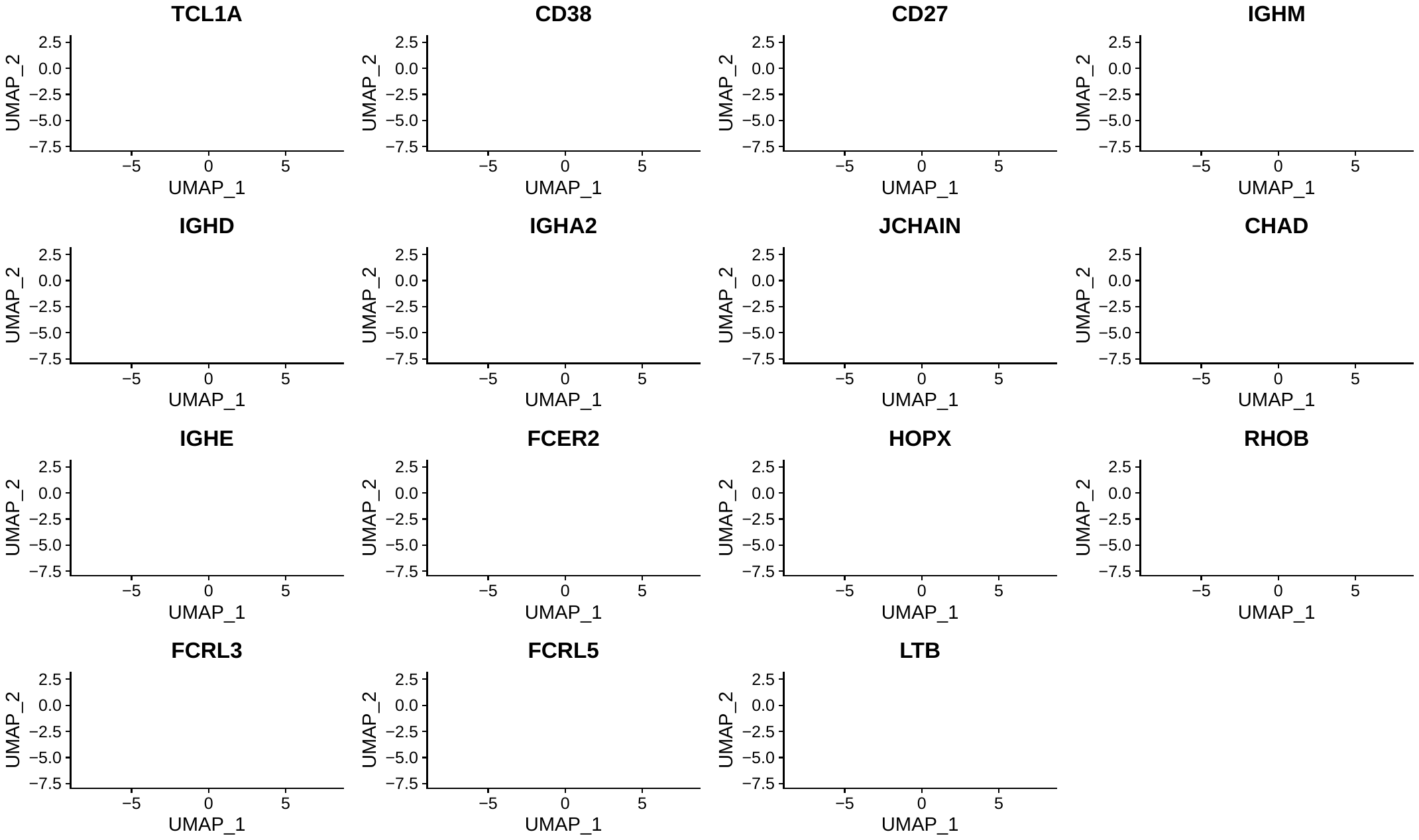

Expression of transcriptomic markers defined in this dataset. Cells are coloured by cluster identities (ref main text Figure 1a) and expression levels are represented by colour intensity.
